# “Decadal climate-driven decoupling between gross primary productivity and tree growth in Mediterranean forests”

**DOI:** 10.64898/2026.02.23.707372

**Authors:** Daniela Dalmonech, Elia Vangi, Dánnell Quesada-Chacón, Alessio Collalti

## Abstract

Mediterranean forests are becoming increasingly vulnerable under climate change, as the growing frequency and intensity of droughts and heatwaves amplify physiological stress, reduce productivity, and heighten the risk of large-scale disturbances. Yet vegetation activity trends, as revealed by remote sensing, may obscure divergent responses between photosynthetic activity and growth, a critical early warning of forest vulnerability. Therefore, the long-term relationship between photosynthesis and tree growth remains poorly understood at regional scales, especially in Mediterranean areas. To address this challenge, we applied a mechanistic, process-based forest ecosystem model across approximately 2,400 km² of Mediterranean forests in southern Italy, encompassing a heterogeneous landscape characterized by diverse stand structures and species dominance. This framework enabled us to explicitly trace carbon fluxes from gross primary productivity (GPP) through allocation processes to average tree growth. By mean of a factorial approach, we identify over extended areas an emergent spatial pattern of divergence of summer GPP and radial tree growth amplified in space and time by the climate variability of the last two decades and shaped by forest legacy. Our findings reveal also that canopy-level greening can mask structural vulnerability and previsual decline across Mediterranean forests. Data show as an apparent long-term trend in photosynthesis decline during summer, not necessarily translates to tree growth decline. Improving our ability to determine if, where and when a key change in forest behaviour will occurs, remains essential for designing effective restoration measure and anticipating tipping points in forest resilience under accelerating climate change.

## 1 INTRODUCTION

Over recent decades, global forests have exhibited a progressive decline in their capacity to function as effective carbon sinks (Forzieri et al., 2022; Pan et al., 2024). This weakening of the forest carbon sink arises from the combined effects of direct anthropogenic disturbances, such as forest harvesting and biomass removal, and an increasing frequency and intensity of change-driven natural disturbances, including droughts, heatwaves, storms, and pest outbreaks (Knutzen et al., 2025; Migliavacca et al., 2025). Since 2000, large areas of European forests have experienced more frequent and severe extreme events, particularly heat waves, and droughts (Lhotka and Kyselý, 2022; Ossó et al., 2022; Xu et al., 2020). Understanding how recurrent extreme events alter carbon fluxes, and the extent to which forest responses mediate these changes, is crucial for monitoring and assessing forest ecosystem functioning. Such knowledge is essential for designing and implementing effective restoration strategies in vulnerable forests, ensuring the recovery and conservation of forest carbon sinks, and thereby supporting the European Union’s climate change adaptation and biodiversity objectives (Chapman et al., 2025; Svensson et al., 2025).

Long-term trends in warming and decreasing precipitation level, coupled with increasing atmospheric dryness, have threatened forest health and growth across Europe and other regions (Samaniego et al., 2018; Sanginés de Cárcer et al., 2018; Yuan et al., 2019; Martinex del Castillo et al., 2022). In southern Europe, the anthropogenic climate signal is becoming increasingly evident (IPCC AR6), with extremes intensifying in both frequency and magnitude. The Mediterranean basin, recognized as a climate change hotspot, is particularly vulnerable: drought strongly affects the socio-economy (Noce et al., 2016; Tramblay et al., 2020) and forest functioning, especially where tree species live at their southern distribution or rear-edge limits (Noce et al., 2017). Although Mediterranean tree species are often considered drought-adapted (Joffre et al., 2007), predicted higher warming rates with increased frequency and intensity of extreme weather events under future climate scenarios are expected to worsen drought-induced dieback, tree defoliation, and impair tree growth (Gazol and Camarero, 2022; Molina et al., 2020; Petritan et al., 2021; Ripullone et al., 2020). Relationships have been demonstrated between lower growth rates, i.e., decline in basal area increment (BAI), and tree mortality risk (Cailleret et al., 2017; DeSoto et al., 2020; Vayreda et al., 2012), potentially predisposing forests to severe drought impacts (Neycken et al., 2024).

Tree growth responses to environmental variability arise from the complex interplay and covariation of multiple abiotic and endogenous biotic factors. Species-specific functional traits, eco-physiological characteristics, and carbon allocation strategies (Decarsin et al., 2024), together with edaphic conditions (Gessler et al., 2018; Sterck et al., 2024) regulate tree growth dynamic. In addition, forest structure and demographic processes (Astigarraga et al., 2025; Wang et al., 2024) shape trees’ capacity to withstand stress and to recover following stressinduced growth declines. Estimates of temporal trends in tree ring data have already proven effective for interpreting growth rate changes in relation to climate change (Piovesan et al., 2008; Schurman et al., 2019; Shestakova et al., 2016). However, dendrometric monitoring is expensive, time-consuming, and spatially sparse, thereby limiting the capacity to assess drought impacts consistently across Mediterranean regions (Tramblay et al., 2020).

Remote sensing data can partially address vegetation monitoring challenges at large scales, over long time periods, and at high resolution (Bathiany et al., 2024). Long satellite records can provide decadal canopy-level assessments, revealing for instance large-scale and widespread greening, modulated by land use and land cover change, increasing atmospheric CO_2_ concentration and temperature but down-regulated by vapor pressure deficit (Zhu et al., 2016; Yuan et al., 2019; Liu et al., 2023). Yet canopy-level greening can mask localized mortality events and disturbance (e.g. Yan et al., 2024) and tree decline may precede detectable canopy-level signal (Neycken et al., 2022; van der Maaten et al., 2024). Vegetation may respond negatively and promptly to extreme conditions such as summer drought, typically captured at very fine scales in the form of local canopy-level damage, while vegetation state might remain largely unaffected as plants can sustain quasi-full recovery at the canopy level (e.g., Italiano et al., 2023; Pollastrini et al., 2019). Hence, climate change and extremes can impact forest ecosystem compartments, e.g. canopy, living biomass, differently and with varying degrees of delay (Kannenberg et al., 2020), therefore posing challenges for monitoring and forecasting near-future forest states and potentially tipping point in forest resilience. To overcome the limitations of dendroecological and remote sensing approach, we use a state-of-the-science biogeochemical, biophysical, process-based forest model to simulate long-term forest dynamics in a Mediterranean region at both canopy and tree levels, including wood formation (Friend et al., 2019). The use of a mechanistic approach to study co-occurring trends of gross primary productivity (GPP) and tree growth has been shown to be promising (e.g., Gea-Izquierdo et al., 2017; Puchi et al., 2026). Simulated forest carbon dynamics and overall stand-level functioning emerge from the internal feedback and interactions among individual biotic and abiotic processes that are mechanistically represented within the forest models. In addition stand-level forest models have been developed for impact studies at higher resolution than their regionallyor globally-resolved counterparts (e.g., ISIMIP-forestry sector; Reyer et al., 2015), to explicitly consider forest structure and management to make predictions (Collalti et al., 2018; Dalmonech et al., 2022; Saponaro et al., 2025). In this study, we apply the process-based forest ecosystem model 3DCMCC-FEM at the regional scale across representative Mediterranean forest ecosystems in southern Italy (Basilicata region). Vegetation activity is quantified through GPP and stand average BAI, capturing both carbon assimilation and biomass accumulation processes. The simulations are conducted over quasi two decades with the specific aim of disentangling and evaluating the relative contributions of carbon assimilation and woody biomass allocation to tree growth. These processes are examined in the context of their modulation by interacting abiotic drivers, such as climate, and biotic drivers, including stand structure and species dominance. Specifically, we address two questions: 1. through which mechanistic pathways does climate forcing influence carbon assimilation, allocation, and growth, and how do these interactions determine long-term tree dynamics in Mediterranean forests?; 2. How do forest structure and species shape the climate induced response of carbon sink and source? Limitations and challenges are also discussed.

## 2. MATERIAL AND METHODS

### 2.1 Climatological data and atmospheric CO_2_

The climate data used in this study were obtained from the ERA5-Land dataset (Muñoz-Sabater et al., 2021), which is a high-resolution (0.1° horizontal resolution) land-surface reanalysis produced by the European Centre for Medium-Range Weather Forecasts (ECMWF). ERA5Land provides hourly data on various land-surface variables, including temperature, precipitation, and surface pressure, among others. Variables not directly available in ERA5Land were first derived from existing data at hourly temporal resolution. Specific (huss) and relative humidity (hurs) were derived using near surface dewpoint temperature, surface pressure (ps), and near-surface air temperature (tas), based on the equations outlined by Buck (1981) and further detailed in Buck (2010). Wind speed (sfcWind) was derived from eastward near-surface wind (uas) and northward near-surface wind (vas) according to sfcWind = sqrt(uas * uas + vas * vas). Sea level pressure over land is computed from the orography (orog), ps, and tas according to psl = ps * exp((g * orog) / (r * tas)), where g is gravity, and *r* is the specific gas constant of dry air. The hourly data was then aggregated to daily values, including minimum (tasmin), maximum (tasmax), mean (tas, ps, psl, hurs, huss, sfcWind), and daily cumulative totals for precipitation (pr), snow fall (prsn), surface downwelling longwave radiation (rlds) and surface downwelling shortwave radiation (rsds), similarly to Lange et al., (2021).

In this study we used tasmax, tasmin, pr, hurs, and rsds for the 1980-2023 period and data were resampled over the simulation grid at 1 km^2^ by mean of a bilinear interpolation.

Global annual atmospheric CO_2_ concentrations were compiled the NASA-observatory dataset (https://scripps.ucsd.edu/) for the period 1980-2023.

### 2.2 Study Area

The region of study, Basilicata, is in Southern Italy and is characterized by a typical Mediterranean climate, with hot summers and mild winters (Figure 1a), and is mainly humid in the Apennine mountains and dry over the hills and flat areas. Elevation gradient ranges from sea level to 2200 meters above sea level, and the precipitation gradient spans from 500 to 2000 mm per year. Forest cover accounts for 29% of the region and represents a broad range of geographical and geological variability, forest species and structure, capturing thus the heterogeneity typical of the forests in the Mediterranean basin (Geri et al., 2010; Vangi et al., 2025). Forests are dominated mainly by deciduous *Quercus ssp* (63%), principally *Quercus cerris* (L.), primarily found in the hilly areas within the elevation belt 500-1000 m, *Fagus sylvatica* (L.)(∼9%), covers mostly areas above 900 m elevation; it follows *Quercus ilex* (L.), (3.4%) and the less abundant coniferous species such as *Castanea sativa* (Mill.), *Pinus halepensis* (Mill.) and *Pinus nigra* (J.F. Arnold) (2.5% and 0.9% respectively) and shown in FigureS1. The primary forest structure is high forest or old coppice transitioning to high forest, ∼58% of the forested area, while ∼27% is managed as coppice (Constatini et al., 2013). The present-day climate in the Basilicata region embeds the fingerprints of the main extremes occurred in the last two decades, which also affected other European forests over large areas, such as the heat waves and drought in years 2007, 2017, 2020 and 2022 (e.g. Lhotka and Kyselý, 2022; Rita et al., 2020), as described by the multi-scalar drought index SPEI (Standardized Precipitation Evapotranspiration Index; Vicente-Serrano et al., 2010) time series reported in Figure 1b.

**FIGURE 1.**
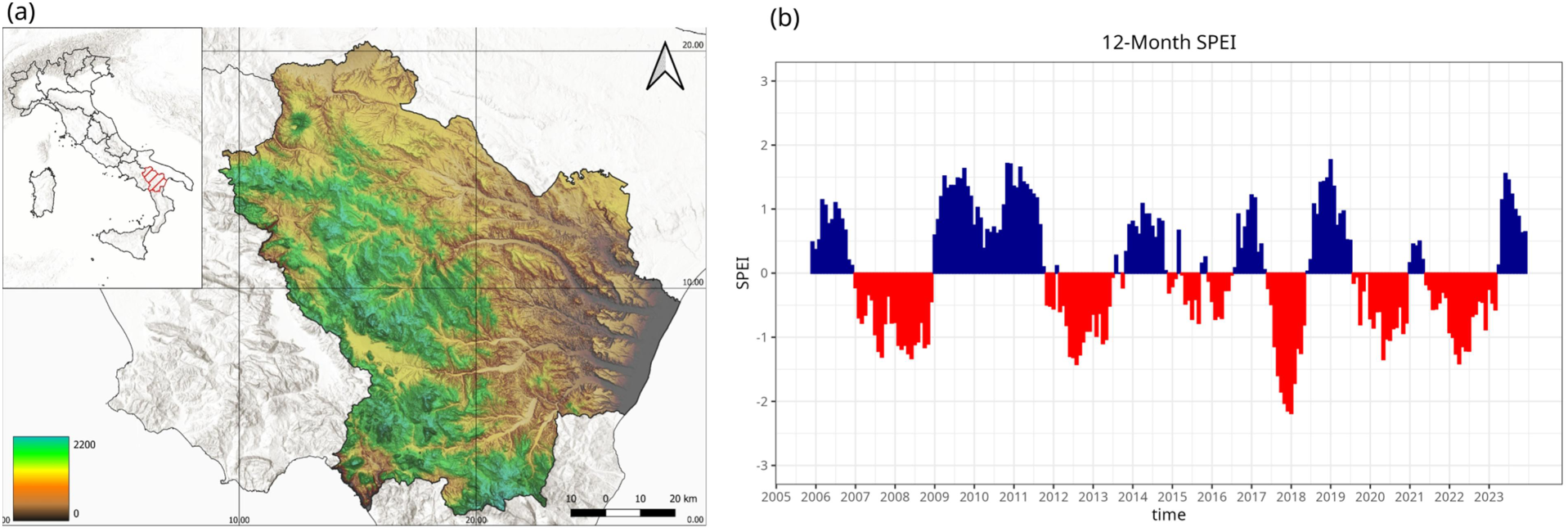
a) Location of the region of study in southern Italy; b) Standardized precipitation evapotranspiration index, SPEI, computed with a temporal scale of 12 months from the ERA5-Land meteorological data. The data represent the median computed across the region; in blue and red, wetter and drier areas than the long-term average, respectively. A SPEI lower than -1 is set as the threshold of onset of mild to severe drought.

### 2.3 The 3D-CMCC-FEM: description and simulation protocol

GPP and BAI were simulated using the process-based forest model 3D-CMCC-FEM (*ThreeDimensional Coupled Model Carbon Cycle – Forest Ecosystem Module*, v.5.6; (Collalti et al., 2024; Collalti et al., 2016; Dalmonech et al., 2022). The model prognostically simulates the carbon and water cycles in forest ecosystems at stand level, explicitly simulating key biochemical and biophysical processes, such as photosynthesis, carbon allocation, and carbon sequestration in woody biomass at species level (Puchi et al., 2026; Saponaro et al., 2025). The model simulates the forest population dynamics with a cohort-like approach, and the eco-physiological dynamics is simulated as responses to environmental drivers and external disturbances such as management. A carbon reserve pool serves as a buffer during periods of negative carbon balance, when plant respiration is larger than GPP and, with a quasi-active role, affects carbon allocation dynamics, i.e allocation of newly assimilated carbon toward stem, leaves or roots. The 3D-CMCC-FEM explicitly accounts for drought impacts, both edaphic and atmospheric, on tree physiology. Direct short-term effects are simulated through the modulation of stomatal conductance using the Jarvis model (Jarvis, 1976), through adjustments in carbon allocation pathways and defoliation. Long term effects arise from temperature-driven increases in metabolic respiration, which reduce plant reserve availability, potentially leading to carbon starvation and, ultimately, tree mortality (McDowell et al., 2008), one of the different ways in the model to simulate mortality. More details on the model are found in section S1 of Supplementary material and Collalti et al. 2024.

The 3D-CMCC-FEM has proven to be a valuable tool for forest ecosystem monitoring (Dalmonech et al., 2024; Vangi et al., 2025), for advancing process-level understanding of forest functioning (Collalti et al., 2020; Puchi et al., 2026; Saponaro et al., 2025), and for predictive applications under changing climatic conditions (Collalti et al., 2018; Dalmonech et al., 2022; Vangi et al., 2024). The 3D-CMCC-FEM was extensively tested and evaluated over a broad range of climate and species, both at local, regional (Dalmonech et al., 2024) and the national scale level (Vangi et al., 2025); in particular the model was shown to realistically simulate GPP sensitivity to daily meteorology, and forest structural attributes related to tree level growth (Mahnken et al., 2022; Puchi et al., 2026; Vangi et al., 2025).

The 3D-CMCC-FEM was here applied over the regional grid at a spatial resolution of 1 km². Initial forest structure was derived for year 2005 from the second Italian National Forest Inventory (INFc 2005) and further refined using high-resolution regional maps of forest and land-use cover, as described in Dalmonech et al. (2024). This approach allowed the model to capture spatial heterogeneity in species occurrence, age distribution, and biomass pools, ensuring that stand-level structural variability and ecological characteristics were adequately represented across the simulated landscape. For each simulation grid cell, a representative forest stand was simulated with one cohort: the most dominant species, hereinafter indicated as ‘forest class’, average diameter at breast height (DBH), average stand density and age. The key species considered in this study were European beech (*Fagus sylvatica*), Black pine (*Pinus nigra*), Sweet chestnut (*Castanea sativa*), Turkey oak (*Quercus cerris*), Aleppo pine (*Pinus halepensis*), and Holm oak (*Quercus ilex*). Species-specific model parameterization was taken from Vangi et al. (2025), while for more details on initial forest structure estimation we refer readers to Dalmonech et al. (2024). Soil characteristics, namely soil texture and soil depth, were extracted from a detailed regional pedological map and the Italian soil map CREA-db (Constantini and Dazzi, 2013). Following the model simulation protocol in Dalmonech et al. (2024), a prescribed constant percentage of the annual thinning rate was applied in forested areas outside protected areas, and areas affected by fire in the simulation period were excluded from the analyses. Overall, the modelled area encompassed approximately 2,400 km².

The 3D-CMCC-FEM was forced by daily data of maximum and minimum temperatures, precipitation, relative humidity and short wave down welling radiation and by global atmospheric CO_2_ concentration (section 2.1). Simulations were performed from 2005, first year of forest structural data availability, until 2023 as “present-day” scenario. For comparison, an additional run was performed, termed “baseline”, terms which will be used throughout this study. The “baseline” forcing scenario was created by randomly sampling years from the 1980-2004 record for both ERA5-land meteorological and atmospheric CO_2_ data as done in (Collalti et al., 2018).

The BAI was used as a proxy for average tree-level growth, partially reducing the age- and size-related effects (Biondi and Qeadan, 2008). Specifically, the BAI was computed from the simulated average DBH assuming a circular shape of the stem, as follows:

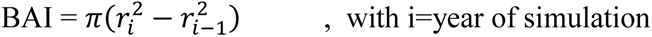

### 2.4 Remote sensing-based data

Selected vegetation indices, from remote sensing (RS), were calculated from the Landsat images bands at a 30-meter resolution and include the (i) Enhanced Vegetation Index, EVI, (ii) Normalized Difference Vegetation Index, NDVI, (iii) tasselled cap greenness, TCG (Crist and Cicone, 1984; Xue and Su, 2017; Jiang et al., 2008).

Those vegetation indexes represent the RS analysis standards and are the most leveraged and widely recognized as standard metrics in remote sensing analyses due to their ability to provide consistent, quantitative information on vegetation status and dynamics (Xue et al., 2017).

In particular, the NDVI delineates vegetation and vegetative stress by showing dense vegetation with high positive values, soil with low positive values, and water with negative values. The EVI index is known for not saturating as rapidly as NDVI in dense vegetation and has been shown to be highly correlated with photosynthesis, plant transpiration, and vegetation biomass in several studies. Finally, the TCG reflects the combined effects of intense absorption in the visible wavelengths caused by plant pigments and high reflectance in the near-infrared, resulting from internal leaf structure and the associated scattering of near infrared radiation, NRI. This spectral pattern is typical of healthy green vegetation. TCG has been shown to correlate moderately-to-strongly with percent canopy closure, leaf area index, and fresh biomass.

Monthly image composites for the study area were generated by selecting the optimal observations for each pixel from all available Landsat-5 TM, Landsat-7 ETM+, and Landsat-8 OLI imagery using the Best Available Pixel (BAP) method (White et al., 2014). This method allows filling the final image mosaic with the best pixel according to four different scoring criteria: (i) acquisition day of the year (DOY), (ii) opacity, (iii) distance to clouds and cloud shadows, and (iv) sensor type. All scores were then summed to provide a total score for each pixel, and the pixel with the highest score (i.e., the BAP) was used in the final image composite. Images acquired within ±15 days of the target DOY with less than 50% cloud cover are considered viable candidates for compositing. As the target DOY, we select the 15^th^ day of the month.

The data were made freely accessible via the Google Earth Engine from the United States Geological Survey (USGS; https://earthengine.google.com). Further information on the BAP image compositing approach can be found in Griffiths et al. (2013), and detailed information on tuning parameters can be found in White et al. (2014).

We additionally selected the GOSIF-GPP dataset (Li and Xiao, 2019 a), which provides global estimates of monthly GPP fluxes at 5 km resolution from solar-induced fluorescence (SIF) data retrieved by the orbiting carbon observatory satellite OCO-2 (Li and Xiao, 2019b). The SIF is the best proxy of plants’ photosynthetic activity estimated over large areas to date (Sun et al. 2018). All RS-based data were masked according to the forest mask and resampled to the 1 km^2^ simulation grid.

### 2.5 Decadal GPP-BAI analyses

Long-term trends (2005-2023) were computed for RS-based data and simulated GPP data, and the occurrence of positive and negative trends, was compared. The Theil–Sen estimator was applied to estimate the slope magnitude and the significance of the slope was assessed using the Mann-Kendall test, accounting for autocorrelation in the time series (Yue et al., 2002). Vegetation activity trends in both RS and modelled GPP were normalized to the average over the simulation period (2005-2023) and reported as percentages. Trends were computed from aggregated values in the summer period, i.e., June, July, and August, which is the most robust signal across RS-based data and models, as it captures the peak of the growing season and thus has a high signal-to-noise ratio (Leisenheimer et al., 2024). Additionally, confounding signals from understory vegetation (e.g., grass, evergreen vegetation) are reduced. The same trend analyses were performed on the modelled annual BAI data.

To determine where and to what extent climate variability, affected long term trends of GPP and BAI in the last two decades, the results under the present-day and “baseline scenario were compared.

Finally, we created a meta-model of the spatially simulated BAI trends at regional scales using a random forests model. This statistical model employs a machine learning technique, based on regression trees, that uses an ensemble of multiple individual random trees (Breiman 2001). This algorithm was chosen for its robustness to collinearity among predictors and the ability to detect linear, nonlinear, and interacting effects among predictors. The algorithm was applied to the simulated BAI trend, training the statistical model with a list of selected spatial predictors.

The selection of variables followed a parsimonious yet ecologically meaningful approach. Key climate predictors included the 2005–2023 spring and summer averages of maximum temperature, precipitation, and vapor pressure deficit, along with GPP and the 12-month SPEI as an aggregated drought measure (Vicente-Serrano et al., 2010). Abiotic predictors comprised soil depth, elevation, and soil clay percentage. Initial stand density, average DBH, and age class were also included as variables related to forest structure.

The final random forest model selection and cross-validation were carried out using the % of explained data variance metric (R^2^). The importance of the predictors and their significance were assessed by the increase in mean squared error (MSE) after variable permutation (Strobl et al., 2007).

For post-processing and statistical analyses, the R programming language (R Core Team, 2021) and the packages *RandomForest* and *rfPermute* (Liaw and Wiener, 2002) were used.

## 3 RESULTS

### 3.1 Model simulations and comparison to satellite data

Normalized time series of GPP during summer as simulated by the 3D-CMCC-FEM and time series of RS-based vegetation activity are reported in Figure 2a along to linear trend and significance of the slope. Interannual variability of vegetation activity well correlates among datasets. It follows the pattern of climate variability depicted by the SPEI_12_ index in Figure 1b, with apparent drops in activity in 2012, 2017 and 2022. Although generally consistent with RSbased observations, the 3D-CMCC-FEM exhibits larger negative anomalies compared to RS data. As result, among long term linear trends, the trend of modelled GPP is positive, yet not significant, as opposite to the RS-based trends. Figures 2b-f show maps of normalized modelled trends of GPP along with the RS-based estimates. The simulated GPP and RS-based vegetation activity show a consistent and coherent spatial pattern characterized by the occurrence of positive trends in the southern-west part of the region (principally covered by Q*uercus cerris* and *uercus ilex*, Figure S1), with lower values, or even negative trends, at the highest elevations, above 1000-1200 m, largely dominated by beech stands (Figure S1). From west to east, moving from the Apennine range to the plain and drier areas, decadal trend values decrease in absolute terms. Among the different RS-based records, spatial modelled GPP trends mirror the pattern estimated from the EVI and GOSIF-GPP time series. Differently from EVI and GOSIF-GPP records, TCG and NDVI-based maps show systematically positive trends in the entire region (Figures. 2d-e and Figure 2f).

**FIGURE 2.**
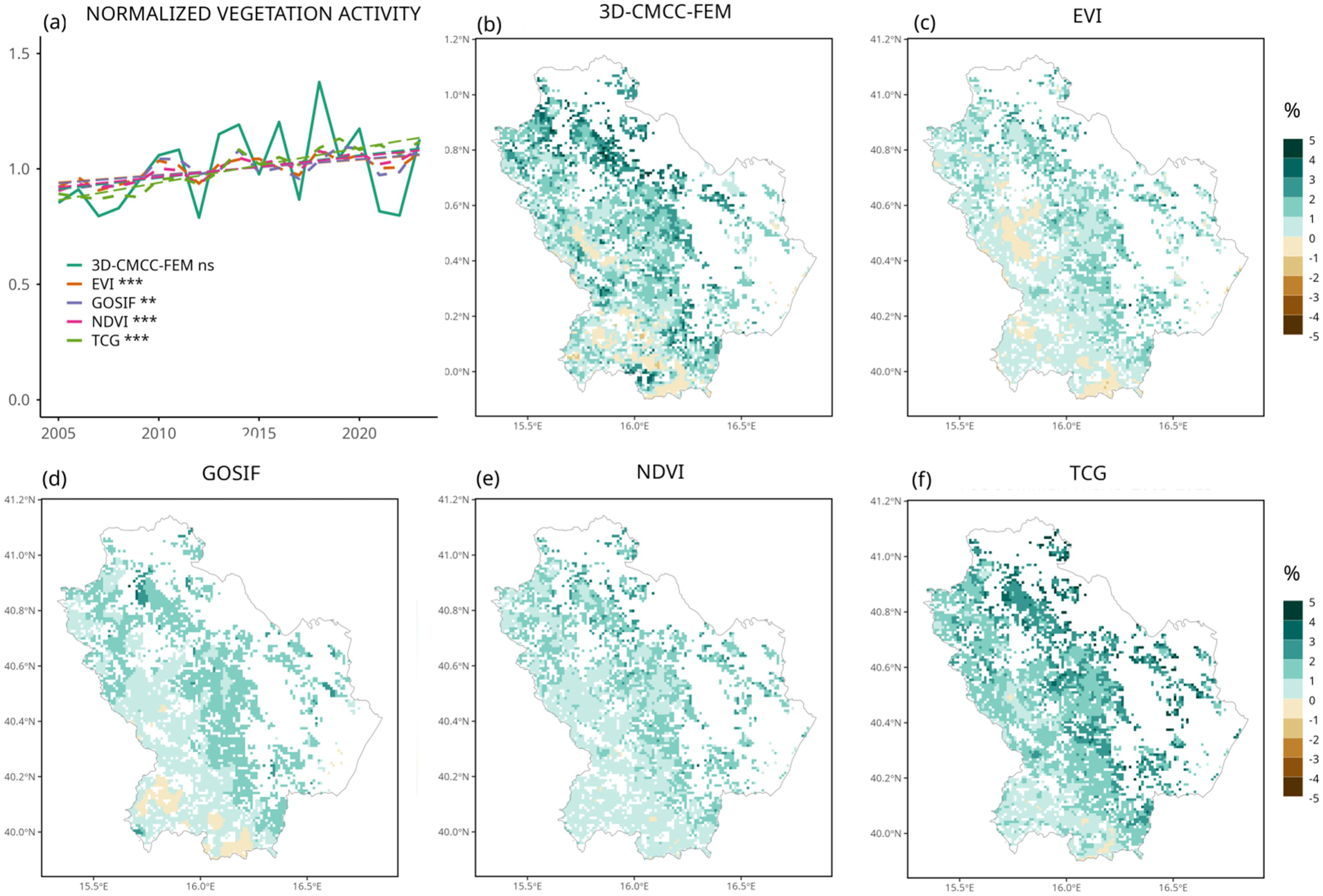
a) Time series of normalized vegetation activities during the summer months (June to August), linear trend slope significance is reports as *ns*, not significant, ** P<=0.01,*** P<0.001; Maps of model and RS-based trends of vegetation activity in summer in the period 2005-2023 for a) modelled GPP in the present-day run, the vegetation indexes b) EVI, d) NDVI, e) TCG, and c) the GOSIF-GPP data. Trends are reported as % change over the mean. White grid cells are areas without forests (according to the INF definition), areas not simulated by the model due to a lack of species parameterization, and areas affected by fires.

When significance of the trend is considered, trends values provided by RS-data, i.e. EVI and GOSIF-GPP, are significant in at least half of the considered simulation grid cells, where positive trends occur (Figure 3a and S3). The modelled trends are instead mostly positive, but not significant, over large areas dominated by deciduous oaks, chestnuts and Aleppo pine (Figure 3b). The trends based on the EVI-record indicating grid cells with statistically significant negative trends, are localized at the highest elevation over the Apennine range, covered by high forests of beech (Figure S1a). While the model also shows negative trends that align with EVI and GOSIF-GPP estimates, these are not, however, significant (Figure 3a-b and S3). Larger differences between EVI and GOSIF-GPP are instead observed when considering the signal for the month march until august (Figure S4).

**FIGURE 3.**
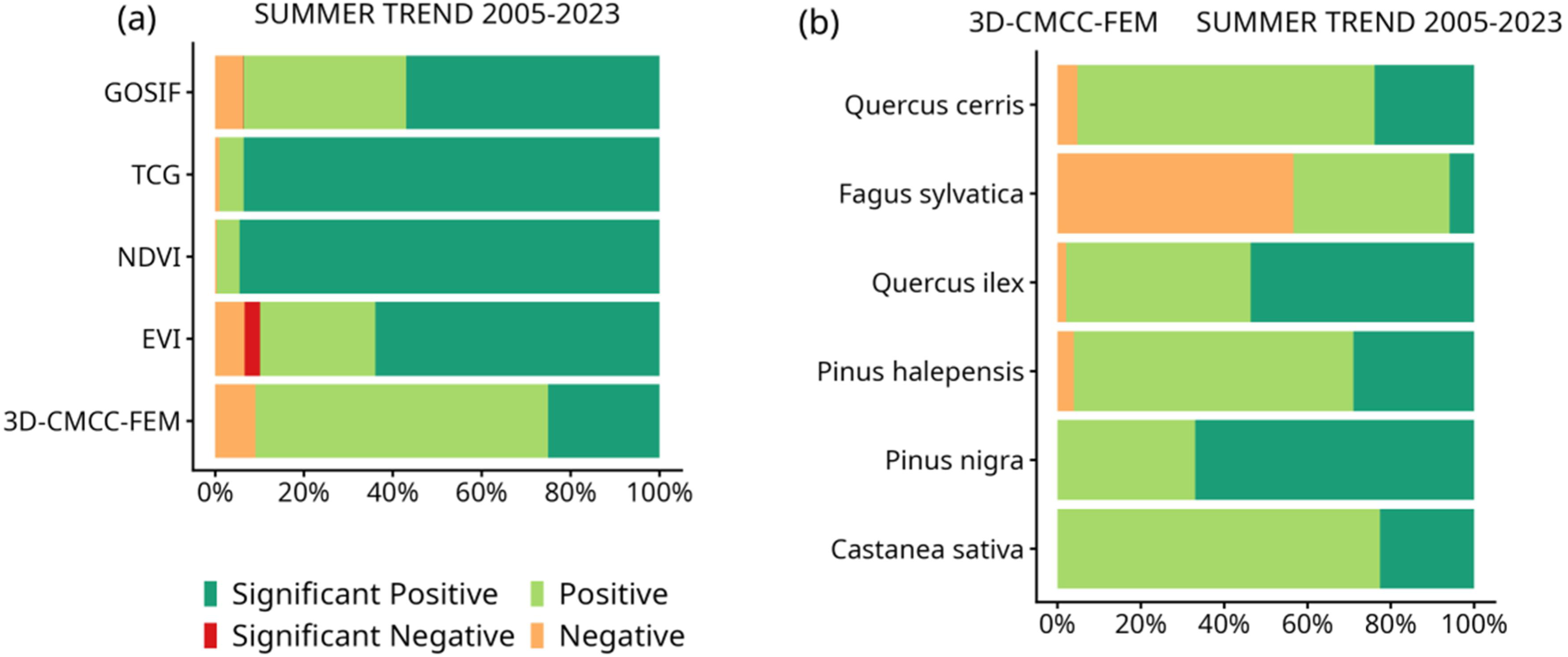
a) Occurrence of positive and negative trends in summer, as a percentage of the total number of the simulation grid cell. Trends are computed for modelled GPP in the present-day run and RS-based vegetation activity data. Significance is considered at P<0.05; b) occurrence of positive and negative trends in the modeled GPP, according to forest class.

When modelled GPP and GOSIF-GPP estimates are considered in more detail, representing the same variable, the two datasets show comparable variability (see section S3 and FigureS5) and satisfactory model performance in terms of spatial patterns and across different temporal scales, i.e. seasonal and interannual (Table S1). Compared to a previous regional application (Dalmonech et al., 2024), where the same forest model was forced with a different meteorological driver, performance generally improved, likely due to a more accurate representation of the precipitation field in the meteorological input.

Model performances at inter-annual time scales show regionally average correlation values of 0.68 and 0.56 for annual and summer anomalies respectively (Table S1); average fractional variance values, i.e., normalized difference in variance between model and estimates of the GPP anomalies, are closer to 1. Modelled Carbon Use Efficiency values, being CUE = 1 – (Plant respiration/GPP), vary between 0.3 and 0.45 (Figure S1c), well within the range provided in literature from observations and ground data estimates (Collalti et al., 2020; A. Collalti and Prentice, 2019; Luo et al., 2025), ensuring that the forest model is well balanced in terms of carbon assimilation and biomass respiration (and net primary productivity). Simulated multiyear average GPP values and BAI values reflect the aboveground forest structures and productivity, with the highest GPP values at the highest elevation dominated by mature beech forest and lower at the lower elevation and toward the plain areas mostly covered by Mediterranean pines (Figures. S1a-b, S2). Higher BAI values are simulated for mature and oldgrowth forests, such as those dominated by large chestnut trees (∼25 cm²), followed by black pine plantations (∼40 cm²). Smaller BAI values are found in the sub-Apennine areas, where deciduous oaks are mostly young stands with coppice structure (∼16 cm²) and beech-dominated high forests (∼7 cm²).

### 3.2 Decadal climate impact on the GPP-BAI trends

The absolute differences in simulated trends of GPP in summer and BAI between the present-day and baseline scenario are reported in Figure 4a and 4b, respectively. The spatial patterns of long-term trend differences are similar for both carbon assimilation and tree growth. Differences are negative in the plain areas, below 200 m, and in the south-western part of the region, particularly within the 500-1000 m elevation belt. In the latter case, compared to the entire region, trends are generally larger under the baseline scenario, i.e. larger negative differences between runs.

**FIGURE 4.**
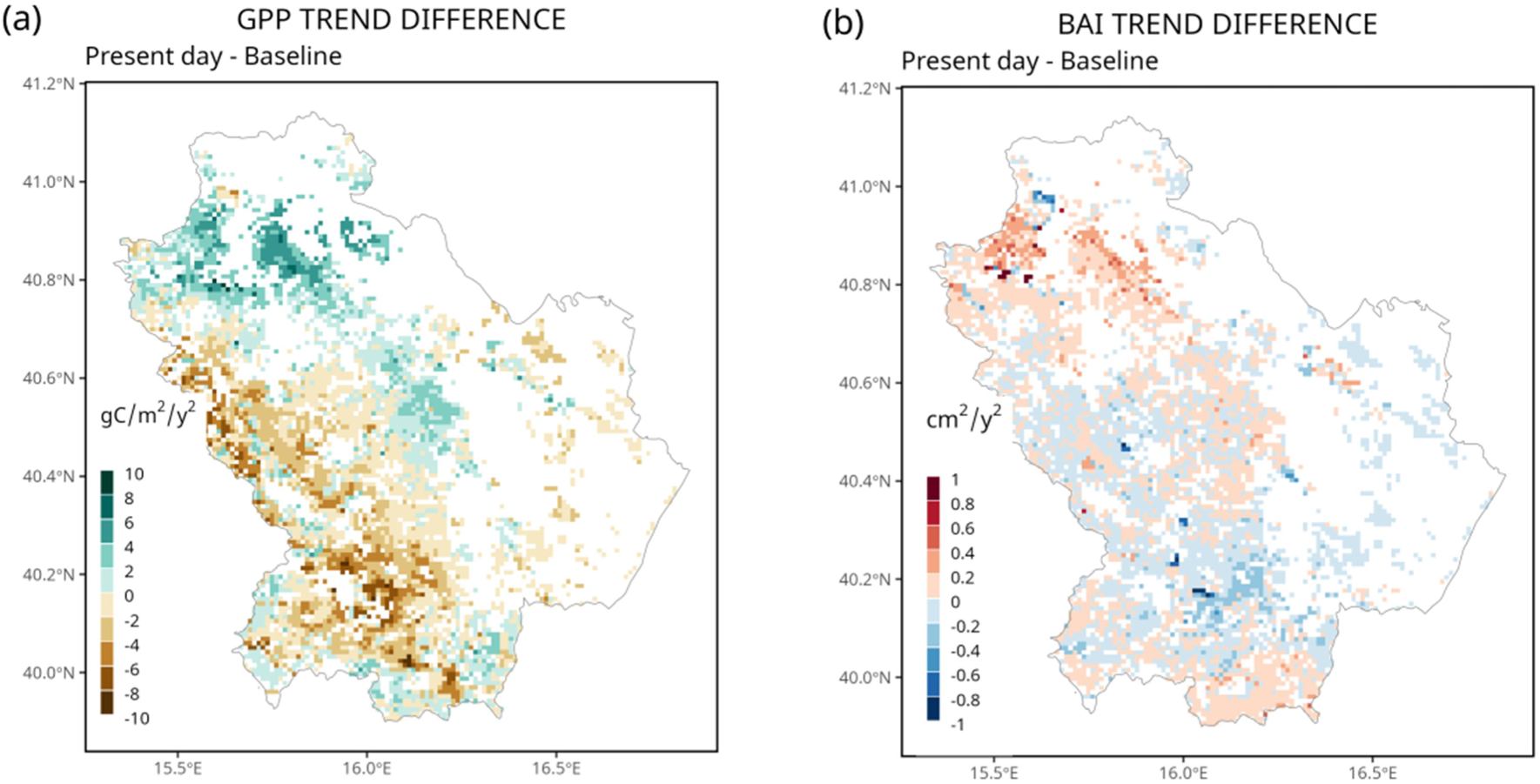
Absolute differences in summer GPP and BAI trends for the 2005-2023 period between present-day and baseline scenario. White grid cells are areas without forests (according to the INF definition), areas not simulated by the model due to a lack of species parameterization, and areas affected by fires.

The sign agreement between trends of GPP and BAI is highlighted by aggregating results into four classes labelled as GPP^pos^-BAI^pos^, GPP^pos^-BAI^neg^, GPP^neg^-BAI^pos^, and GPP^neg^-BAI^neg^ where superscripts indicate the sign of the trend (Figure 5a and 5b). Under the baseline scenario, and across most of the simulated areas, growth patterns closely mirrored those of GPP (Figure 5a), highlighting a strong coupling between carbon assimilation and biomass accumulation. In the southern-western part of the region, in the middle-to-low-elevation forest belt, GPP trends are mainly positive, while BAI trends are negative (Figure 5a). The change in forested area falling into each GPP-BAI trend classes in the present-day scenario compared the baseline scenario are reported in Table 1.

**FIGURE 5.**
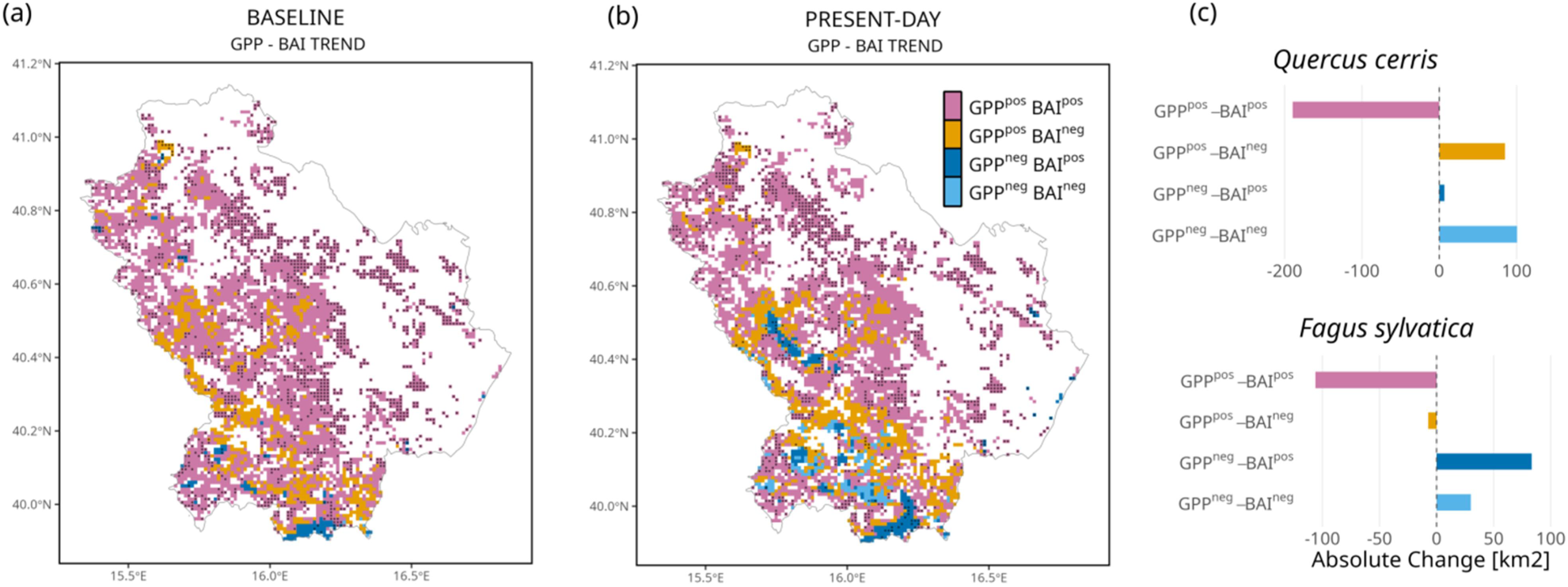
Occurrence of positive and negative trends of summer GPP and BAI according to sign association under a) baseline and b) present-day scenario. Areas where BAI trends are significant (P<0.05) are indicated by hatching. White grid cells are areas without forests (according to the INFc definition), areas not simulated by the model due a to lack of species parameterization, and areas that have been affected by fires. c) Net change between present-day and baseline scenario of forest cover within the GPP-BAI class reported for areas dominated by *Q. cerris* and *F. sylvatica*.

**Table 1.**
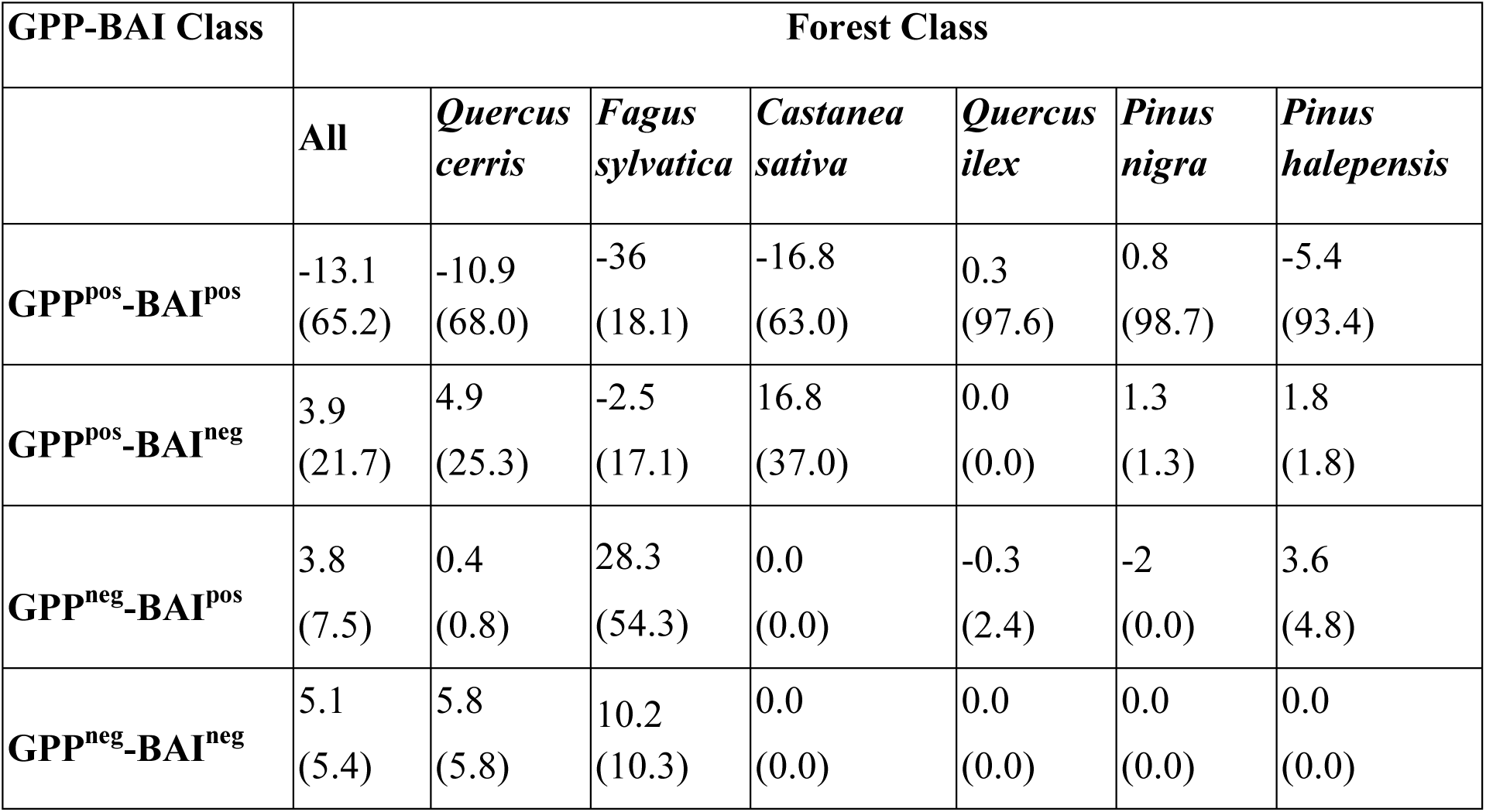
Forest cover change in each GPP-BAI association class between present-day and baseline scenario. Data are reported as percentage, according to forest class, i.e. forested areas dominated by a single species. GPP data refer to summer months, June to August. Within brackets, for each forest class, the percentage of forest cover falling in each GPP-BAI association class under the present-day scenario is reported.

| GPP-BAI Class | Forest Class |  |  |  |  |  |  |
| --- | --- | --- | --- | --- | --- | --- | --- |
|  | All | *Quercus*<br>*cerris* | *Fagus*<br>*sylvatica* | *Castanea*<br>*sativa* | *Quercus*<br>*ilex* | *Pinus*<br>*nigra* | *Pinus*<br>*halepensis* |
| **GPP<sup>pos</sup>-BAI<sup>pos</sup>** | -13.1<br>(65.2) | -10.9<br>(68.0) | -36<br>(18.1) | -16.8<br>(63.0) | 0.3<br>(97.6) | 0.8<br>(98.7) | -5.4<br>(93.4) |
| **GPP<sup>pos</sup>-BAI<sup>neg</sup>** | 3.9<br>(21.7) | 4.9<br>(25.3) | -2.5<br>(17.1) | 16.8<br>(37.0) | 0.0<br>(0.0) | 1.3<br>(1.3) | 1.8<br>(1.8) |
| **GPP<sup>neg</sup>-BAI<sup>pos</sup>** | 3.8<br>(7.5) | 0.4<br>(0.8) | 28.3<br>(54.3) | 0.0<br>(0.0) | -0.3<br>(2.4) | -2<br>(0.0) | 3.6<br>(4.8) |
| **GPP<sup>neg</sup>-BAI<sup>neg</sup>** | 5.1<br>(5.4) | 5.8<br>(5.8) | 10.2<br>(10.3) | 0.0<br>(0.0) | 0.0<br>(0.0) | 0.0<br>(0.0) | 0.0<br>(0.0) |

Under the present-day scenario, the percentage of forest in the class GPP^pos^-BAI^pos^, is ∼65% of the total cover, with a decrease of -13.1% compared to the baseline run (Table 1). The change is mirrored by a corresponding net increase in forested areas in the classes GPP^pos^-BAI^neg^, GPP^neg^-BAI^pos^ and GPP^neg^-BAI**^neg^**, i.e. +3.9 %, +3.8 % +5.1% respectively (Table 1). Modelled results indicate how the forest cover characterized by positive trends of GPP and negative trends of BAI under present-day conditions is ∼21%. A similar qualitative result is also observed when considering the GPP signal during the growing season (Figure S6), with a net increase in forested areas described by opposite GPP-BAI trends.

When considering forest classes, about 68% of deciduous oak forests currently falls into the class GPP^pos^-BAI^pos^ (Table1), and 25.3% in the class GPP^pos^-BAI^neg^.The latter mostly located in the low hilly bioclimatic belt up to about 1000 m (Figure S7). Compared to the baseline scenario, deciduous oak forests show a net loss of area in the class GPP^pos^-BAI^pos^ of -10.9%, corresponding to ∼ 190 km^2.^.The net change is mirrored by an increase of a +4.9% in the GPP^pos^-BAI^neg^ class, ∼ 85 km^2^, (Figure 5c), and +5.8% increase of areas in the GPP^neg^-BAI^neg^ class, ∼ 100 km^2^ (Figure 5c Table 1).

In beech dominated forests under present-day scenario, the model shows as only 18.1 % of areas are characterized by positive trends of both GPP and BAI, while the majority, 54.3 % belong to the class GPP^neg^-BAI^pos^ principally occurring at high elevations, in the range of 1100-1800 m (Table 1 and Figure S8) and 17.1% to the class GPP^pos^-BAI^neg-^. The remaining beech forests, namely 10.2%, are in the class GPP^neg^-BAI**^neg^** and are mostly located in the low-elevation range (Figure S8). Similarly to oak dominated areas, beech forests also show a net change of area from the class GPP^pos^-BAI^pos^, -36% corresponding to ∼ 100 km^2.^(Figure 5c). This change principally reflects an increase of areas described by negative summer GPP with positive trends in BAI, +28.3% corresponding to ∼ 83 km^2^ (Figure 5c).

The results for the other forest classes, that altogether cover about 6.9% of the total forested area of the region, are only reported in Table 1. Model results indicate relatively minor changes in the frequency of GPP–BAI classes. Holm oak and black pine exhibit net changes of approximately 1%, while Aleppo pine shows a net change of ∼5% in the GPP^pos^-BAI^pos^ class. However, only a few and sporadic grid cells are affected. Chestnut forests display a more pronounced decrease, with a 16.5% reduction in the GPP^pos^-BAI^pos^ association class. While the four GPP-BAI classes shown in Figure 5 are spatially robustly localized, the significance of the negative trends for the long-term trend BAI is very sporadic, yet it occurs in beech-dominated forests and in the north-eastern part of the region.

### 3.2 BAI trend spatial pattern

The Random Forest algorithm was used to rank the importance of the selected predictors in statistically explaining the simulated spatial pattern of BAI trends. The model embedding structural variables, soil-related variables, and using SPEI as the climate-related driver alone explained about 73% of the BAI trend variability (Figure 6 inset). All the predictors are statistically significant in explaining the spatial variability of BAI, although with varying degree of sensitivities. Initial stand density was the most important explanatory factor, followed by age class, GPP, average SPEI_12_ values during spring, and average initial DBH (Figure 6). No notable differences were observed when meteorological variables were explicitly included (Figure S9). In the latter case, stand density is still the most important explanatory variable; however, more relevance in the ranking is given to the precipitation signal during summer and soil depth, yet most of the predictors have a corresponding increase of MSE in the Random forest model below 30%, indicating the challenges of the statistical model to discriminate variable importance robustly.

**FIGURE. 6.**
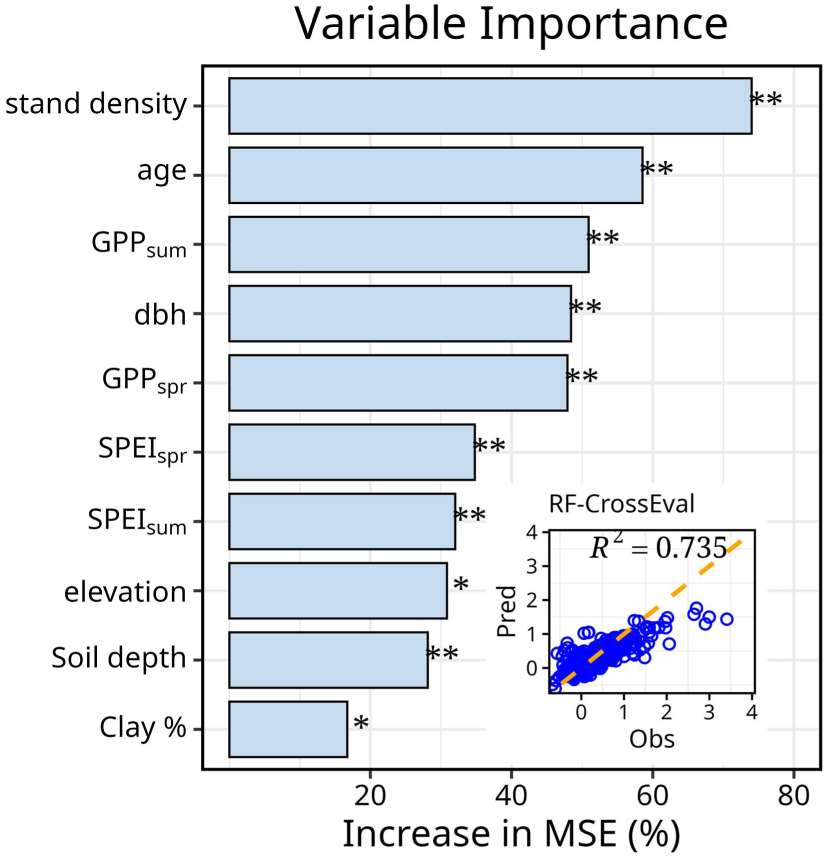
Variables importance in explaining spatial patterns of BAI trends according to the random forest model (RF) algorithm (number of trees=1000, training on the 70% of the data, cross-evaluation on the 30%). Importance is reported as increase in the mean squared error, MSE, with permutation testing for significance (* P<=0.05** P<0.01). Subscript indicated *sum*, summer, *spr*, spring. Inset: cross-evaluation result of the random forest model, R^2^ coefficient of determination.

## 4. DISCUSSION

Climate change impacts do not affect all components of forest ecosystems uniformly, nor do they occur synchronously, with responses often differing in magnitude and timing among different ecosystem compartments (Kannenberg et al., 2020), particularly in Mediterranean regions (e.g. Nolè et al., 2013). In this study, we investigate the long-term effects of climate variability over the past two decades on forest functioning in a Mediterranean area of southern Italy by means of a factorial approach to separate biological trends. This study provides an ecosystem level and broad-scale perspective on how key processes, such as seasonal carbon assimilation and tree growth, have responded to changing climatic conditions, highlighting potential shifts in productivity patterns, forest resilience, and ecosystem carbon dynamics

### 4.1 How Do Climate Influence decadal Carbon Processes and Forest tree dynamics in Mediterranean forests?

Modelled results indicate that recurrent droughts over the past two decades have exerted a sustained negative effect on both seasonal carbon assimilation trends and tree growth, with growth trends likely being more strongly impacted. Despite the negative climate impact, both the 3D-CMCC-FEM results and independent RS-datasets show a coherent spatial pattern, characterized by a general summer greening over most of the region under study, mostly significant for observations. The observed positive trends in GPP are consistent with the “CO₂ fertilization effect” hypothesis (Gonsamo et al., 2021; Zhu et al., 2016), which is explicitly represented in the forest model. This hypothesis suggests that rising atmospheric CO₂ concentrations can enhance plant photosynthesis, potentially stimulating forest gross productivity. At the same time, forest management practices contribute to landscape heterogeneity, further modulating these productivity patterns. Indeed, forest inventory data estimates show the occurrence of relatively young forest in the Mediterranean compared to e.g. central Europe, a condition arising from both natural and anthropogenic drivers, with large areas managed as coppice (INF 2021). The high frequency of stand in young age classes, typically indicated to show a higher CO₂ fertilization effects (e.g. Collalti et al., 2018; Norby 2016) and the natural stand development before fully canopy closure, likely contribute to the observed positive vegetation activity trends. Under these conditions, increases in growth, are very well known to be associated with positive trends following stand development, hence trends of carbon assimilation and tree-level growth are expected to align. Yet, it has also been shown, how even mature forest with large trees might still show positive BAI trends (e.g. Fekedulegn et al., 2003; Stephenson et al., 2014).

Interestingly, our results also highlight an emerging large-scale pattern, where specific areas in the region show a divergent trend in seasonal carbon assimilation and growth, divergence which is already occurring under the baseline scenarios, yet amplified under present-day climate, with more forested areas affected by this divergence. The rate of uncoupling between carbon assimilation and growth remains a longstanding topic of debate (Boukhris et al., 2025).

Scientific evidence indicates that photosynthesis and diametric growth can become decoupled over short (monthly) to annual timescales (Anderson-Teixeira and Kannenberg, 2022; Cabon et al., 2022; Martínez-Sancho et al., 2022) when fuelled by carbon reserve. Other studies have shown that, at annual scales, tree-ring chronologies and remote sensing-based proxies of vegetation activity tend to be closely correlated across forests in Spain and southern Italy, although the strength of these associations may vary with species and edaphic conditions (Castellaneta et al., 2022; Vicente-Serrano et al., 2020). Differently, the present study reveals a diverging pattern of assimilation and growth emerging at decadal scales, which is supported by local observations of growth decline in deciduous oak sites. Following recent droughts and heat waves, several deciduous oak-dominated stands in the regions have experienced dieback, with healthy and weakened trees coexisting within the same site (Colangelo et al., 2017; Conte et al., 2025).

Previous studies suggested that dry conditions might explain the uncoupling of the C-source and -sink, i.e., the tree-ring with a GPP-based signal observed on-site (e.g. Cabon et al., 2022). In this context, during the summer, stomatal closure may limit carbon assimilation by trees (Gessler et al., 2018). Nevertheless, carbon reserves generally remain sufficient to sustain metabolic functions, albeit at the expense of stem growth, which in turn can influence carbon allocation patterns in subsequent years (Huang et al., 2021). This imbalance of the carbon allocation, as regulated actively by the carbon reserve pool (Gessler and Zweifel, 2024), might loop back on longer temporal scales and under frequent drought events. Trees rely on reserves to recover from hydraulic damage, such as embolism, and to maintain metabolic functions under stress (Hammond et al., 2022). However, if unfavourable conditions persist, repeated or prolonged reserve consumption can trigger long-term negative feedback: reduced carbon availability limits stem growth and other allocation pathways (e.g., to leaves), weakening tree vigour. Peltier et al. (2024) demonstrated that trees in drier sites in the US experience elevated Non-Structural Carbohydrate, i.e. carbon reserve, consumption and reduced mixing in the sapwood under prolonged dry conditions, because photosynthesis can stop but not respiration. This cascade can ultimately lower stand-level resilience and reduce carbon use efficiency, and trees become increasingly vulnerable to further and subsequent stress events (Collalti et al., 2020; He et al., 2020; Peltier et al., 2023). The observed modelled uncoupling between seasonal carbon and average tree growth, while general in the reported mechanistic explanation, it is not however systematic over the entire Mediterranean region under study and other diverging trajectories are shown, with patterns that are also shaped by other biotic and abiotic factors, beside the external climate forcing as indicated in the next section.

### 4.2 How forest structure and species shape the climate induced divergence of sink and source?

Our results show how a general greening over the entire region under study, might mask a negative trend of tree growth, a precursor for the loss of resilience capacity, hence in turn increased vulnerability and tree mortality risk (Cailleret et al., 2017; Neycken et al., 2022; Pedersen, 1998). This occurrence of negative and diverging trends holds particularly for deciduous oaks and beech dominated forests at the lower elevations. Across the other forest classes the positive trends of both carbon assimilation and tree growth in forests dominated by holm oaks reflects the capacity of several evergreen oak species to withstand periods of drought (Heinrich et al., 2025). Similarly, among pine species, Aleppo pine has been shown to have high plasticity and drought tolerance resulting in a wide geographical distribution in the Mediterranean, although a reduced growth was observed in the drier areas of its distribution following severe drought events (Dorman et al., 2015; Gazol et al., 2017).

Notably, the model shows a tendency for GPP and growth trends to diverge, even under the baseline scenarios, indicating that site legacy, primarily reflected in the forest structure estimated for 2005, plays a critical role in driving forest dynamics trajectories under a changing climate (Astigarraga et al., 2025; Petritan et al., 2021). Incorporating current ground-based forest observations, such as stand density as model initial conditions, enables the representation of growth legacy (Kannenberg et al., 2020; Vayreda et al., 2012). Accounting for these legacy effects has proven critical for accurately assessing drought-induced mortality risk (Sterck et al., 2024). For instance, highly dense forests show slower growth compared to less dense forests, a trend that can be exacerbated under drought conditions and soil moisture scarcity (RubioCuadrado et al., 2018).

Besides stand density, age and size of the average tree modulate the forest’s sensitivity to climate variability. Higher-elevation forests are dominated by older age classes and larger trees, with individual growth remaining positive, as also recently observed in the Balkan region (Markuljaková et al., 2026). Although larger trees are theoretically and empirically associated with higher respiration costs (Collalti et al., 2020), their size also corresponds to greater carbon reserve pools. These observations align with recent studies indicating that old forests can, under current climate conditions, maintain high resistance and potentially resilience under both Mediterranean and temperate climates (Colangelo et al., 2021; Puchi et al., 2024). This holds for instance for European beech, a diffuse ring species, which shows no negative BAI trends at higher elevations despite negative trends in the summer signal of carbon assimilation and vegetation activity. Larger size of these mature stands and ring characteristics might have led to larger pools of carbon reserve and the ability to buffer year-to-year climate anomalies impacts more efficiently. However, the tree ability to buffer climate extremes impact depends not only by the size but also by its physiology traits and edaphic conditions, making the impact of drought and extremes not univocal.

At medium to high elevation, beech trees might be more energy limited and thus additionally benefit from an increased carbon assimilation rate during warmer springs or earlier snowmelt (Nezval et al., 2025; Signarbieux et al., 2017), which might lead in turn to an earlier start of wood enlargement before the summer heat-wave (Grossiord et al., 2022). In turn, the modelled tree growth trend appears unaffected by the negative summer GPP trends indicated by both the model and several RS products. At lower elevations, beech stands tend to show a growth decline. Enhanced vegetation activity during the spring might likely amplify the drought impact as transpiration occurs at higher rates, potentially decreasing soil water availability later into the year (Bastos et al., 2021), contributing to the drop in carbon assimilation and the end of the spring season, and in turn reserve consumption and growth (Bose et al., 2025). This pattern, differently from results in Martinex del Castillo et al. (2022), shows thus a much more heterogeneous response for beech dominated forests.

Generally, at national level has been shown how in the period 2010-2022, beech-dominated sites are undergoing an increasing rate of defoliation, which was, however, not accompanied by a significant increase in tree mortality; conversely deciduous oaks, i.e., Q*uercus cerris*, showed a significant increase in mortality rates (ICP-forest monitoring network plots; Bussotti et al., 2024).

Compared to beech, deciduous oaks have a porous ring, hence, a small amount of active tree ring and sapwood. Therefore, oaks might respond more tightly to the current-year drought event with long-term impacts (Zweifel and Sterck, 2018). In oak forests in the dry-subhumid areas of the study region, a persistent combination of low water resource availability and declining growth may deplete carbon reserves, might hamper the capacity to sustain metabolic processes and repair hydraulic damage, despite oaks being known to be drought-tolerant species. This dynamic can increase tree vulnerability, potentially leading to higher mortality risk and reduced stand resilience under expected increase of recurrent drought events in the near future (Desprez-Loustau et al., 2006; Gentilesca et al., 2017; Piper et al., 2022).

### 4.4 Further considerations and challenges

Despite the robustness of the forest model, some sources of uncertainty in model performance and process representation required further discussion and clarification. In order to keep model simulation feasible and constrain the simulations, a simplified setup was used to initialize the model, which also had to align with the details of the soil and climate inputs, the latter still provided at a relatively coarse resolution. Model simulations do not account for changes in stand species composition, and thus it does not capture the potential occurrence of more drought-tolerant tree species.The model uses physiological, species-specific parameters that are, however, general and not locally calibrated, which might partially explain the lower occurrence of statistically significant trends compared to RS-based data. Similarly the lack of significance in the modelled trends might be the result of the high model responsiveness to summer precipitation shortage, a behaviour previously reported for other models (e.g. 3-PG; Nolè et al., 2013) and for the 3D-CMCC-FEM model and attributed to the parsimonious representation of soil water dynamics (Saponaro et al., 2025).

Other potential extremes affecting Mediterranean forests are missing, such as frost events (D’Andrea et al., 2021). However, the effect of frost is currently not included in the model, and it would require highly accurate daily meteorological data to simulate its impact. Similarly, the model has been developed to run offline at daily time scales, with readily available meteorological drivers. Therefore, it does not simulate sub-daily processes, particularly finer mechanisms of the plant hydraulic system, which might lead to sudden mortality events without growth decline (Gessler et al., 2018). However the 3D-CMCC-FEM rather aims to capture the average stand behaviour and the long-term quasi-active role of carbon reserves in influencing the allocation of newly assimilated carbon, e.g., toward assimilation rather than growth (Gessler and Zweifel, 2024), and despite the aforementioned uncertainties, we show that the model provides consistent, explainable temporal and spatial trends of GPP and BAI. The role played by the initial stand structure is prominent, so that the persistence of GPPgrowth long term trend divergence does not significantly differ in terms of extent and location of affected forested areas if the model is driven with a different forcing dataset (data not shown). However, different climate forcing might shift the timing of divergence and strengthen trends. In this respect, the needed accuracy of the initial forest structure and climate data to constrains future forest dynamics trajectories in time and space, put emphasis on initiatives such as the *Global Ecosystem Dynamics Investigation* (*GEDI,* Duncanson et al., 2022), or climate impact research initiative such as ISIMIP (Frieler et al., 2024), providing high resolution climate data and climate projections.

## CONCLUSIONS

This study provides holistic evidence of Mediterranean forests functioning under recent climate change in southern Italy, accounting for multiple drivers of long-term association and decoupling between carbon assimilation and tree growth, eventually leading to growth decline. We identify how and to what extent decadal climate variability in a Mediterranean region affected trends in carbon assimilation and tree woody growth, pinpointing areas already showing a background growth decline, often masked by long-term canopy-level greening. The reconstructed forest dynamics in representative Mediterranean stands revealed thus emerging large scale and diverse trajectories of carbon assimilation and tree growth, while accounting for structural differences, species-specific eco-physiology, and climate variability impacts over the last two decades.

Even in the absence of an evident, visible, long-term decline in functioning, or conversely, an apparent decline of functioning, continuous monitoring activities in particular by means of mechanistically approaches as shown in this study, are shown to be critical to investigate how canopy level functioning translates to tree growth. Monitoring activity, informed by knowledge of the actual forest structure and, in parallel, accurate climate data, are pivotal for optimizing target management and the effectiveness of restoration measures.

While this work applies to the Mediterranean forest ecosystem, the transient nature of forest dynamics under anthropogenic climate change makes the results more general.

## Acknowledgments

This research was supported by the European Union – NextGenerationEU under the National Recovery and Resilience Plan (NRRP), Mission 4 Component 2 Investment 1.4 Call for tender No. 3138 of December 16, 2021, rectified by Decree n.3175 of December 18, 2021, of the Italian Ministry of University and Research under award Number: Project code CN_00000033, Concession Decree No. 1034 of June 17, 2022 adopted by the Italian Ministry of University and Research, CUP B83C22002930006, Project title “National Biodiversity Future Centre – NBFC”.

## Author contribution

D.D. conceived the study, designed and performed the model simulations. D.D. and E.V-. analyzed the data. E.V. and Q.D. were responsible for data curation. A.C. was responsible for software and funding acquisition. All co-authors contributed to the discussion of the results and the review and editing of the manuscript.

## Data Availability Statement

The 3D-CMCC-FEM model code version 5.6 is publicly available under the GNU General Public Licence v3.0 (GPL) and can be found on the GitHub platform at: https://github.com/Forest-Modelling-Lab/3D-CMCC-FEM). All data, model executable, and scripts to perform analyses and figures presented in this work are provided open access in the Zenodo server (https://doi.org/10.5281/zenodo.18701467). Correspondence and requests for additional materials should be addressed to the corresponding author.

## Notes

https://doi.org/10.5281/zenodo.1870146

### Competing Interest Statement

The authors have declared no competing interest.

